# Computational Engineering of a Therapeutic Antibody to Inhibit Multiple Mutants of HER2 Without Compromising Inhibition of the Canonical HER2

**DOI:** 10.1101/2023.07.21.550003

**Authors:** Sapir Peled, Julia Guez-Haddad, Nevet Zur Biton, Guy Nimrod, Sharon Fischman, Yair Fastman, Yanay Ofran

**Affiliations:** The Mina & Everard Goodman Faculty of Life Sciences, Bar-Ilan University, Israel; Biolojic Design, Ltd., 12 Hamada Street, Rehovot 7670314, Israel

## Abstract

Genomic germline and somatic variations may impact drug binding and even lead to resistance. However, designing a different drug for each mutant may not be feasible. In this study, we identified the most common cancer somatic mutations from the Catalogue of Somatic Mutations in Cancer (COSMIC) that occur in structurally characterized binding sites of approved therapeutic antibodies. We found two HER2 mutations, S310Y and S310F, that substantially compromise binding of Pertuzumab, a widely used therapeutics, and lead to drug resistance. To address these mutations, we designed a multi-specific version of Pertuzumab, that retains original function while also bindings these HER2 variants. This new antibody is stable and inhibits HER3 phosphorylation in a cell-based assay for all three variants, suggesting it can inhibit HER2-HER3 dimerization in patients with any of the variants. This study demonstrates how a small number of carefully selected mutations can add new specificities to an existing antibody without compromising its original function, creating a single therapeutic antibody that targets multiple common variants, making a drug that is not personalized yet its activity may be.

## Introduction

Cells contain naturally occurring germline and somatic variations [1]. Cancer cells have more mutations than healthy cells due to somatic mutations [2]. COSMIC, a database for somatic mutations in cancer (cancer.sanger.ac.uk) [3], currently records over 23 million genomic variants in ∼1.5 million clinical cancer samples, an average of 15 somatic mutations per sample. These somatic and germline mutations may affect the binding of drugs to their targets in different ways, compromising efficacy and conferring resistance [4]. Somatic and germline mutations have been shown to cause resistance to several anti-cancer drugs, *e.g.* EGFR mutations that cause cetuximab or gefitinib resistance [5, 6], Topoisomerase II mutation that causes Amascarine resistance [7] and HER2 mutations that cause resistance to trastuzumab [9] and to several small molecules [8, 10]. According to one account, in nearly half of cancer patients pre-existing resistance to chemotherapy is observed prior to, and independent of, treatment due to existing variations [11]. Thus, for many drugs, response rate is significantly reduced because of mutations that compromise efficacy. Possible effects of such mutations are steric hindrance of the drug binding pocket [12], target protein conformational changes [13], or reduced binding affinity [14]. It has been suggested that a customized design that considers specific variations in the drug target, may lead to new versions of existing drugs that will be more effective in patients with specific mutations [15]. However, mutations in the populations are usually overlooked in drug development [16, 17], and the development of mutation-specific versions of existing drugs poses commercial and regulatory challenges [17, 18].

A possible route for overcoming the challenge of genomic variations that cause drug resistance is the design of multi-specific drugs that can effectively bind common variants of the target protein. A single molecule that can bind slightly different variants of the target can provide a single general treatment that is produced, developed, and regulated once, and yet takes into consideration personal genomic profiles.

Antibodies are increasingly used as cancer therapeutics, due to their high affinity, exquisite specificity, long half-life, and their mammalian and sometimes even human origin [19]. Several formats of engineered antibodies posses the capability of more than one different antigens [20]. Among these formats, dual-specific antibodies were designed to target two distinct targets with a single antibody (“Two-in-one antibody”) [21–25]. Unlike bi-specific antibodies, which have different arms that bind different targets, dual-specific antibodies may be symmetrical IgGs, in which both arms can bind both targets. We propose that such antibodies can be computationally designed to recognize both wild-type and mutated versions of targets. Such a single multi-specific compound may address a wider range of mutants [26]. In infectious disease, a broadly neutralizing antibody (bnAb) is a naturally occurring antibody that binds multiple variants of the same protein, *e.g.*, in attacking the highly variable HIV-1 virus [27]. In the context of infectious diseases, such multi-specific Abs are directed toward one conserved epitope that is assumed to be the same in all variants. In the context of cancer, however, somatic cancer mutations that confer resistance to an effective treatment [28] may not leave a conserved active site that can be attacked by a broadly neutralizing antibody. Thus, inhibiting multiple variants with a single antibody requires the design of a multi-specific antibody that binds slightly altered epitopes to block known escape paths of the tumors [29]. While such an approach may not necessarily block all possible mutants and may create selective pressure for the emergence of new escape mutants, it will nonetheless increase the response rate and extend the duration of the response.

HER2, a human epidermal growth factor receptor (also known as ERBB2), is a known oncogene and is overexpressed in several cancers, *e.g.*, breast [30], ovarian [30], gastric [31], salivary glands [32] and lung [33]. In breast cancer, it is over-expressed in about 15-30% of the tumors and is associated with poor prognosis [34]. HER2 mutations are found in 4% of all breast cancer patients and can go up to 10% in specific breast cancer subtypes. They are abundant in metastatic ER+ tumors, suggesting a role in acquired resistance to endocrine therapy [35]. In COSMIC v.95, there are 941 different cancer somatic genomic variations in HER2. There are four approved anti-HER2 antibodies, and five biosimilars [36]. Pertuzumab (Trade name: Perjeta®)[37] is a cancer treatment, approved for use in combination with Trastuzumab (trade name: Herceptin®)[38] and chemotherapy for treating breast and gastric cancers [39].

Pertuzumab inhibits HER2 dimerization, thus blocking the HER signaling pathway [37]. Here, we analyzed reported somatic mutations in cancer patients and identified two HER2 mutations, S310F and S310Y, that, according to our predictions, may confer resistance to Pertuzumab. Previous studies had shown that Pertuzumab does not bind these mutants [40, 41], although HER2 dimerization is not disrupted by them [41]. In fact, due to *de-novo* hydrophobic interactions [40, 42], these mutations were found to stabilize and enhance HER2 dimerization [41, 43]. Cells expressing these mutations show hyperphosphorylation of HER2 and an increase in the subsequent activation of signaling pathways [40, 42]. Targeting these HER2 mutants is clinically important, as they were found to have oncogenic activity [42] and to promote cell migration, invasion, aggregation, and growth [40, 42, 44]. Each of these mutations has been identified independently as a somatic mutation [45], and the S310Y was obsereved as a germline variation (ClinVar Variation ID: 376188). Thus, targeting them can be relevant not only for patients that were born with these mutations and do not respond to treatment but also for patients that acquire these mutations during their illness and become resistant. While there is no clinical observation that these mutations alone confer resistance to Pertuzumab, it is clear from biochemical studies that Pertuzumab does not bind these mutants. Using computational and experimental methods, we altered the original therapeutic antibody and turned it into a multi-specific antibody. The designed antibody retains its binding to canonical HER2 while also binding the two mutants S310F and S310Y. This computationally guided design of a multi-specific antibody suggests a framework for therapeutic antibody design that may allow for therapeutic antibody candidates that could be used for larger patient populations without requiring the development and regulatory approval of new, personalized therapeutic antibody for smaller patient populations.

## Results

### Therapeutic Abs Target Mutations

COSMIC Mutation Data (Targeted Screens and Genome Screens) was filtered to find mutations in contact sites of therapeutic antibodies. Table 1 shows the number of mutations in each step of the filtration process. We found 102 known missense mutations that alter a residue in the interface between a clinical-stage antibody and its target but do not alter protein length. The most frequent mutations were HER2 S310F and S310Y. These mutations are present on the interface with Pertuzumab (trade name: Perjeta)[37] which targets HER2 for breast cancer treatment. Each of these mutations has been identified independently both as a germline variation (rs1057519816 in dbSNP) and a somatic mutation (COSMIC Mutation Genomic IDs COSV54062198 and COSV54062802, respectively).

**Table 1.** The number of COSMIC mutations after every filtration step, starting from the initial number of mutations with repetitions for different samples.

| Step | Number Of Mutations |
| --- | --- |
| COSMIC Mutation Data | 37,433,225 |
| Mutations in therapeutic abs targets | 220,029 |
| Unique Mutations | 13,561 |
| In contact sites | 740 |
| In FDA approved for cancer | 124 |
| Substitution- missense mutations | 102 |

### HER2 mutations disrupt Pertuzumab Binding

Two HER2 somatic cancer mutations from COSMIC are located in the interface with Pertuzumab as calculated from the 3D structure of the complex (PDB id: 1S78) and are fairly prevalent in the population. Figure 1A shows that the predicted effects of the S310F and S310Y mutations on the Ab-Ag affinity as calculated by Maestro [46] are 3.48 and 2.62 kcal/mol. Note that residue numbering is different between the PDB structure and the Uniprot [47] sequence.

**Figure 1.**
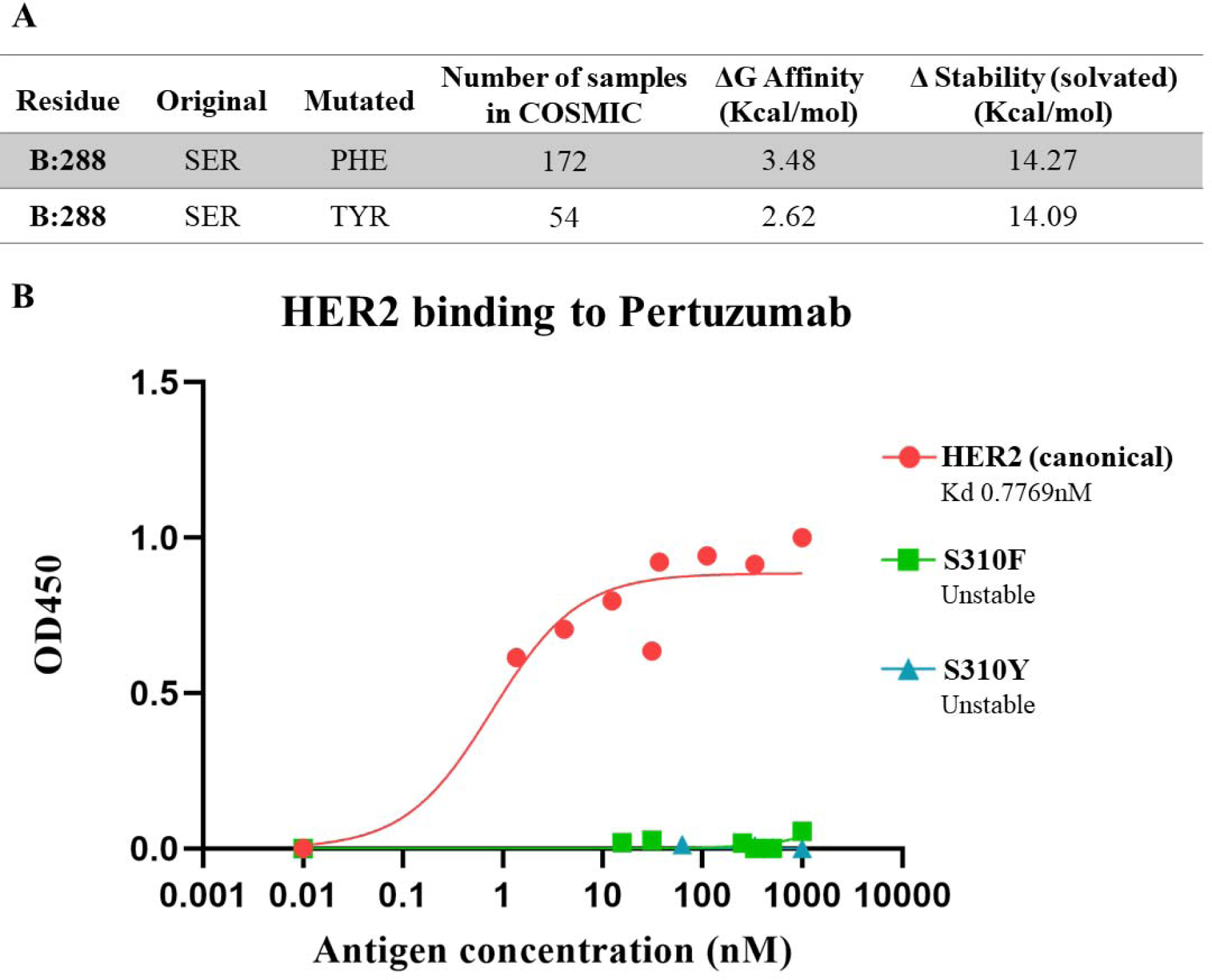
A Mutation analysis results from Maestro. Each line shows a different mutation, with the number of samples in COSMIC containing the mutation, predicted changes in Ab-Ag affinity and Ag stability between the original and mutated states. ΔG Affinity of the S288 mutations is close to or bigger than 3 Kcal/mol, suggesting that they might disrupt HER2 binding by Pertuzumab. S288 in the PDB structure numbering corresponds to S310 in the sequence numbering. B ELISA results for Pertuzumab binding to canonical HER2 and the two HER2 mutants, with Kd values. Both S310 mutations abrogated HER2 binding to Pertuzumab.

S288 in the PDB structure numbering corresponds to S310 in the sequence numbering. Unless otherwise specified, below we will use the Uniprot numbering. Using a previously suggested cutoff [48] of 3 kcal/mol as a guide for a significant predicted change in affinity, we estimated that these mutations will disrupt binding. The complete alignment of the canonical HER2 sequence and the crystalized residues is shown in Supplementary Figure 1. We experimentally validated these predicted effects of the mutations on the therapeutic antibody binding. Figure 1B shows experimentally measured effects of the mutations on Pertuzumab-HER2 binding for canonical HER2 as well as for the two mutants. Both S310 mutations abrogated HER2 binding to Pertuzumab. For our next objective, *i.e*., designing a multi-specific antibody, we attempted to engineer an antibody that binds canonical HER2 as well as both S310 mutants.

### Selections Yielded Tri-Specific Binders

To computationally engineer a variant of Pertuzumab that binds canonical HER2 and its main mutants, S310Y and S310F, we explored two strategies. The first approach was to search for mutations that can strengthen the interaction by directly altering the patch on the antibody surface that contacts S310. The other approach was to try and form new additional contacts in areas of the antibody that are in close spatial proximity to residues in HER2 but do not form energetically beneficial contacts with Pertuzumab. To this end, we designed a library based on Pertuzumab, focusing on antibody residues that are in close spatial proximity to S310. We also included variations in residues that are not in direct contact with HER2 but are near residues that form contacts with the goal of sampling conformational changes in the region that can strengthen binding. To maintain the antibody structure, we avoided residues with multiple intra-molecular bonds and residues that are evolutionarily conserved. This resulted in fourteen positions in which we introduced diversity, allowing all 20 possible amino acids in each position. Figure 2 shows a structure of the Pertuzumab-HER2 complex, highlighting SER310 in cyan (numbered as 288 in the PDB residue numbering) and the fourteen selected Pertuzumab residues for the library, color-coded by the reason for choosing them: Direct contact with S310, spatial proximity to the S310, adjacency to a residue contacting S310. We also added diversity in antibody residues that contact the HER2 in other positions that are not S310, to allow for the strengthening of the complex through alternate interactions.

**Figure 2.**
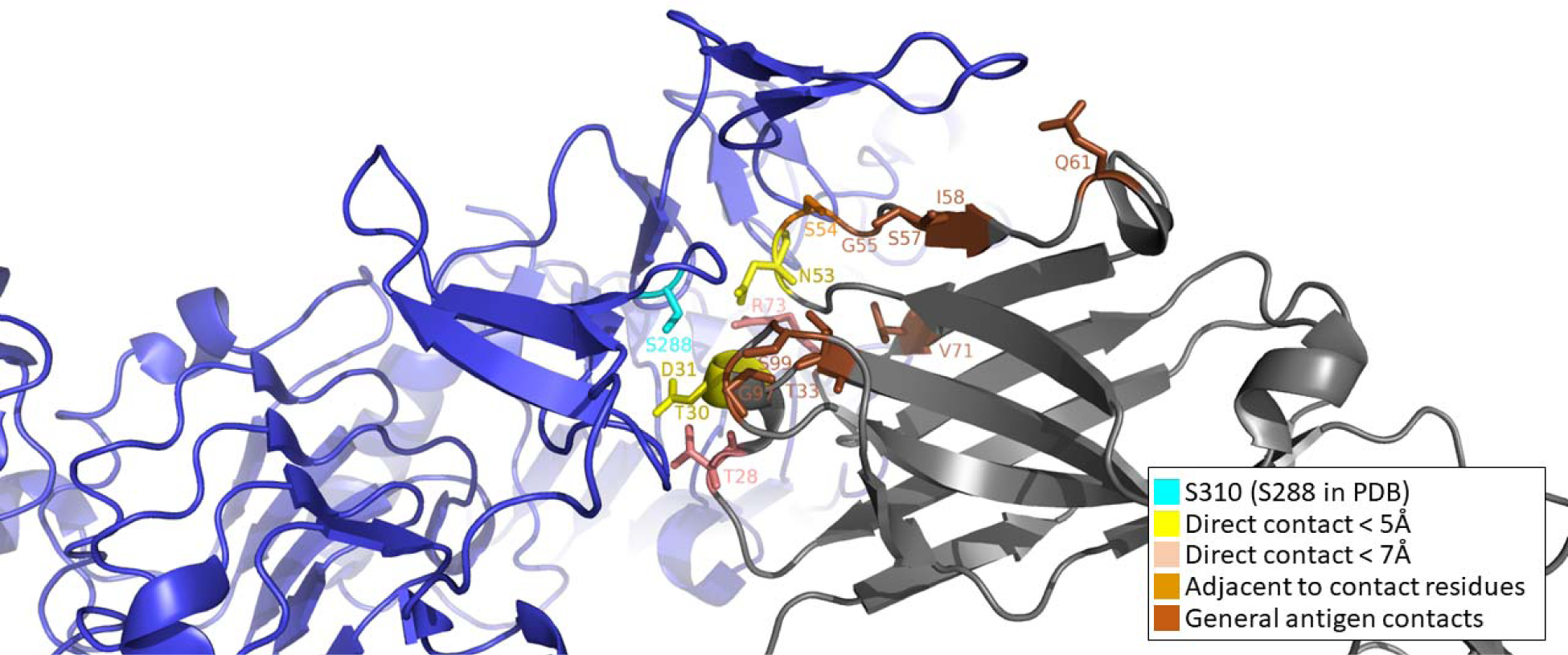
The interface between HER2 and Pertuzumab (PDB 1S78). HER2 is in blue, Pertuzumab light and heavy chains are in light and dark grey, respectively. Serine 310 (288 in the PDB numbering) is shown in cyan. All 14 residues chosen for the library are colored by the criteria that led to their selection for the library.

Figure 3A shows all 14 residues on Pertuzumab heavy chain that were selected for variation, with their corresponding number on the PDB 1S78 structure. Experimental constraints in yeast transformation and yeast surface display limit tractable library size to 10^9^ variants. Hence, with reasonable undersampling, we wished to reduce the diversity of the designed library so that it’s within two orders of magnitude of the experimentally tractableize. We thus limited the number of mutations in each variant such that no more than six mutations are allowed in one sequence in the library. This also helped assure that the variants in the library are not too different from the Pertuzumab sequence and are more likely to maintain a developable profile. The 14 positions selected for the introduction of variation were divided into four groups based on their sequence position, as shown in Figure 3B and the last column of Figure 3A. Based on this division we constructed the library such that each variant in the library will have representative mutations from each group, covering all possible combinations. Figure 3C shows the predetermined number of residues that were mutated from each of the four groups to reduce the diversity of the library, sampling for a total of 3.86×10^11^ sequences. Using yeast surface display the library was selected against the three variants. Figure 4 shows the complete selection scheme, with two MACS rounds and two FACS rounds. After three rounds of selection, a population of putative binders was gated. Figure 5A shows the scatter plot of this gated population against each of the three variants of HER2. As illustrated by the figure, this population appears to contain binders for all three HER2 variants. The cells in the top right quadrant of each plot, which represent putative binders, were collected for the next round of selection.

**Figure 3.**
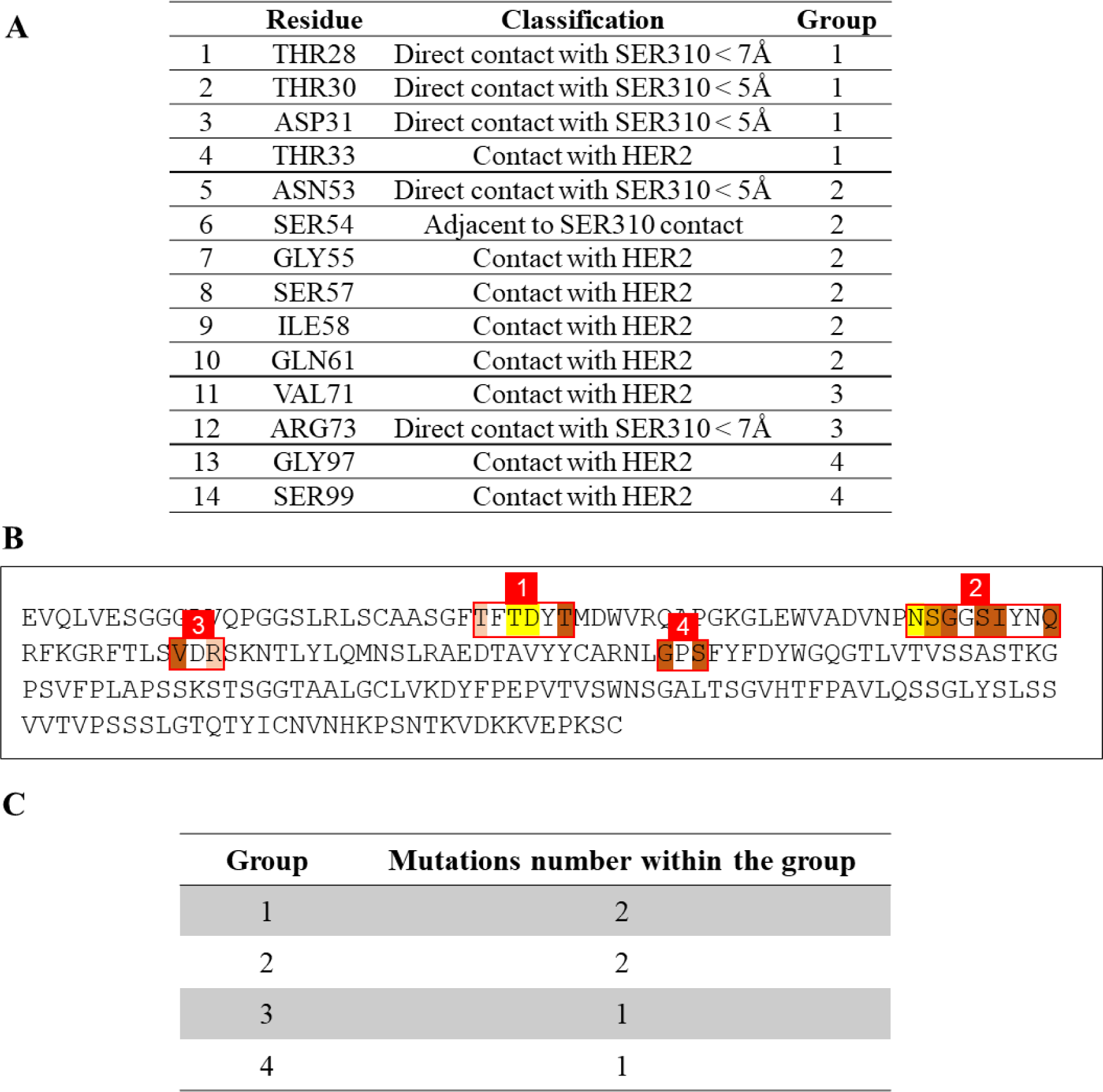
**A** Fourteen putative residues from Pertuzumab heavy chain that were chosen to be mutated. Each residue is classified for the reason it was chosen: in direct contact with SER310, adjacent to a SER310 contact, or a general contact with HER2 that was chosen to strengthen binding. Each residue was assigned to one of four groups based on its sequence position. **B** Pertuzumab heavy chain from the 1S78 PDB structure. Highlighted are the fourteen residues to be mutated, colored by the same color scheme as in Figure 2. The residues are divided into four groups based on their sequence position. **C** The number of residues selected for mutation from each group of residues for a specific library sequence. Groups 1 and 2 which contain four and six residues, respectively, had two mutations each in every library sequence, with all possible combinations of two mutations from each group. Groups 3 and 4, having two residues each, had one mutation in every library sequence. Each library sequence will have a total of six mutations.

**Figure 4.**
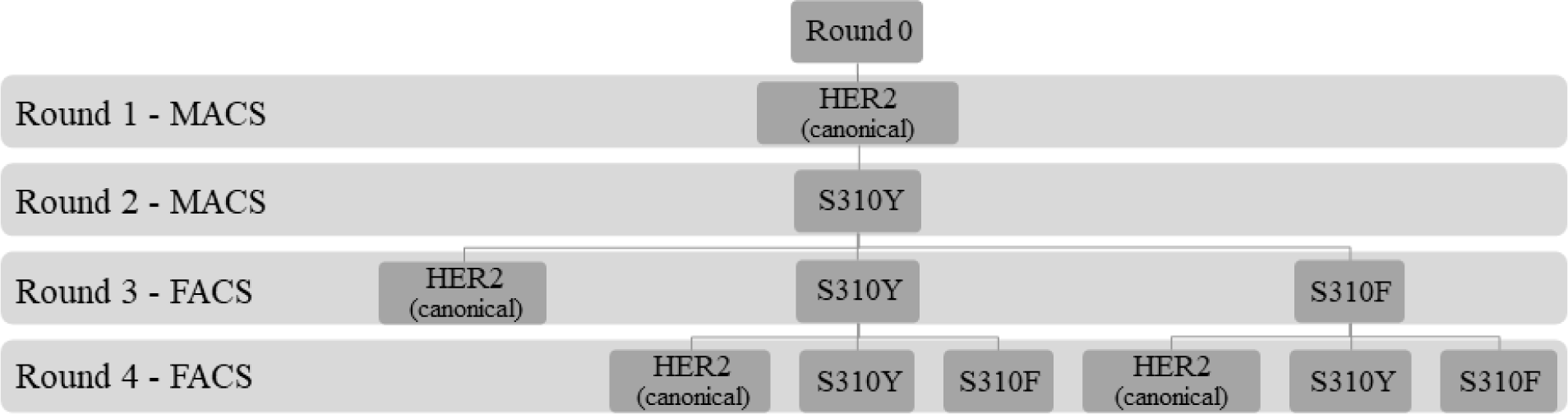
Selections scheme. Two rounds of MACS, one against canonical HER2 and one against the S310Y mutant reduced the size of the initial library. A round of FACS (round 3) was used to gate putative binders that were taken into the fourth and final round of selection, yielding putative binders to be sequenced.

**Figure 5.**
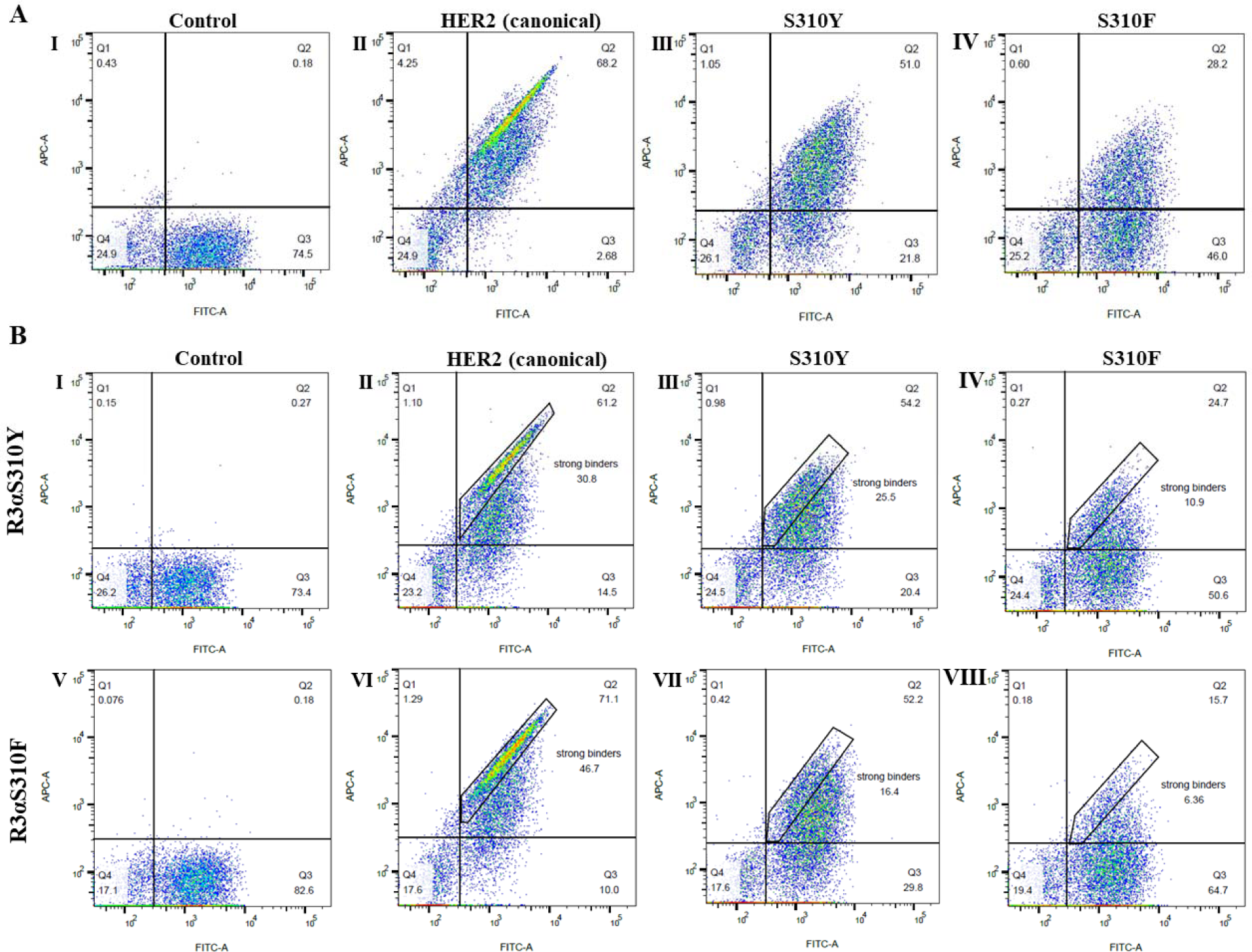
A Binders population after two round of MACS – one against canonical HER2 and the second against S310Y mutant underwent a round of sorting by FACS (R3). Gates were set as follows: I no antigen for control. II 200nM of canonical HER2 III 200nM of S310Y mutant IV 200nM of S310F mutant. The populations from the top right quadrant that represent S310Y and S310F putative binders, were collected for the next round of selection. B The populations from Figure 5A III and IV were taken through an additional selection round (R4). Top panels describe the results of R4 where R3 was selected against the S310Y mutants; bottom panels are R4 where R3 was against the S310F mutants. FACS gates were set as follows: I,V Null, negative control, where no antigen is added, II,VI 50nM of canonical HER2 III,VII 50nM of S310Y mutant IV,VIII 50nM of S310F mutant. Gated populations are marked in rectangles.

### Stringent Library Selection Provided Stronger Binders

Colonies from the third selection round (R3) against both S310Y (Figure 5A –IV) and S310F (Figure 5A –III) mutants were taken through an additional selection round to examine these binders against the second mutation and canonical HER2 in the search for multi-specificity, as shown in Figure 5B.

### Finding Multi-specific Candidates from R4 FACS Results

A total of 90 colonies were sampled, 15 colonies from each antigen-binding population of R4 (Figure 5B: II-IV, VI-VIII). The colonies were sequenced and tested against the three antigens. None of the colonies bound all three antigens. However, three colonies showed binding to two different antigens, as shown in Figure 6A. These three candidate colonies were taken for further analysis as monoclones. The sequences of the 14 mutated positions from all 90 colonies are shown in Supplementary Figure 2.

**Figure 6.**
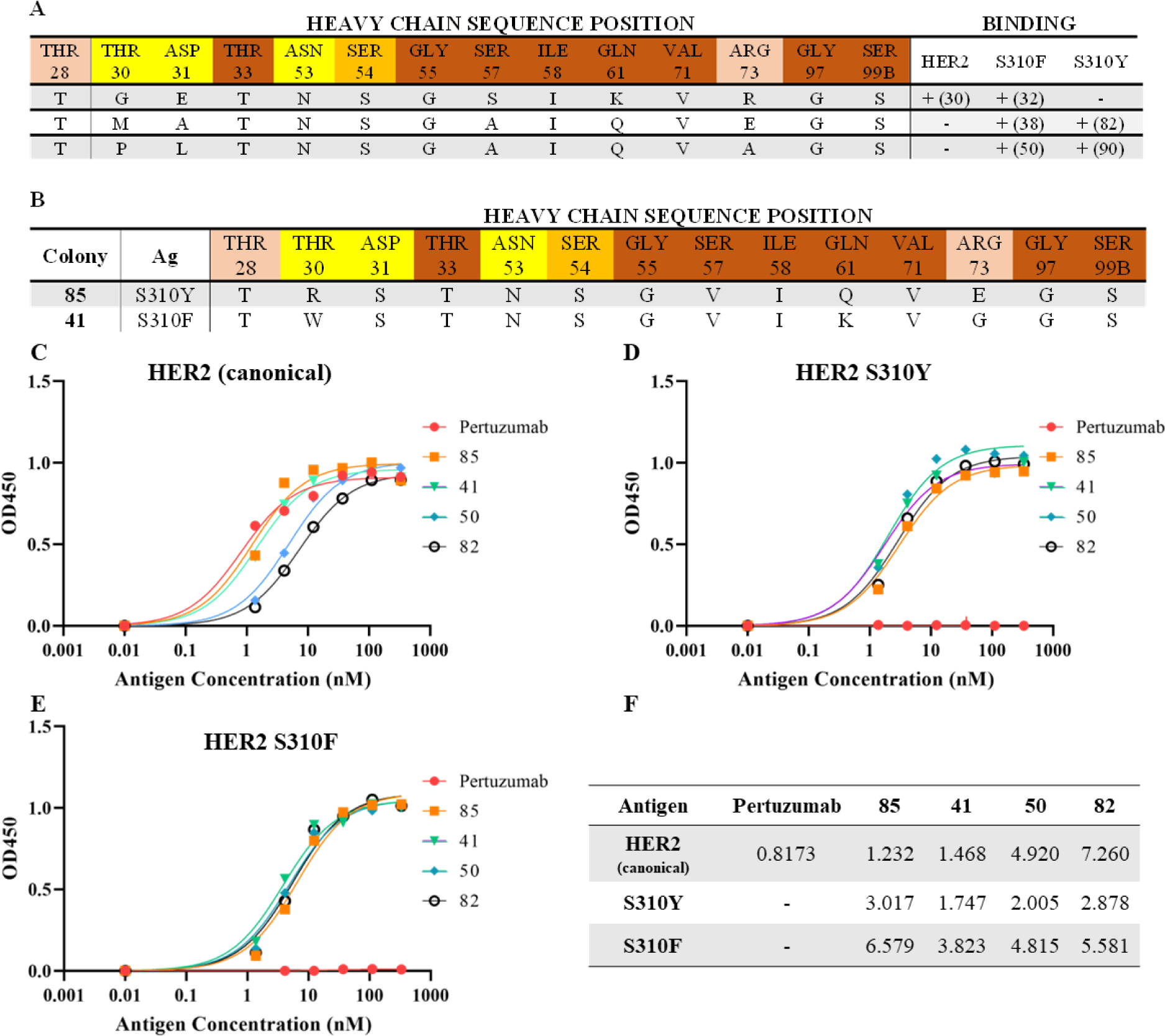
**A,B** Positions and variations that contribute to multi-specific binding according to selection results. The first row of each table shows the fourteen positions from the original Pertuzumab sequence selected for variation, as shown in Figure 3A, numbered as in the PDB 1S78. Positions are colored according to the color scheme in Figure 2: Direct contacts in yellow and peach, contact-adjacent in orange, and general antigen contacts in brown. **A** Three sequences from round 4 of selection, that were found to bind two different variants. On the right side of the table, for each sequence, the antigen it binds is specified. The colony number from which the sequence originated is in parenthesis. **B** Two colonies from round 4 of selection. Each of these sequences is composed of residues that appear in binders of all three antigens: canonical HER2, S310F, and S310Y. For example, according to Supplementary Table 1, in position 58 there can only be Isoleucine or Leucine. Both sequences have Isoleucine in this position. **C,D,E** ELISA results for Pertuzumab and four IgGs from the tested monoclones, against three antigens: **C** canonical HER2 **D** S310Y mutant **E** S310F mutant. **F** Kd values in nM for each IgG described in Figures 3C-E, against all antigens tested. No binding was detected for Pertuzumab and both mutants.

The sequencing analysis did not identify a single sequence that bound all three variants of the HER2. We thus used the sequencing results to try and stitch together such tri-specific sequences. We analyzed the diversity observed in each position and identified all the residues that were observed in all binders of each of the three antigens – canonical HER2, S310Y, and S310F. These residues were then combined, to compose a set of residues per position that appear in at least one binder for all three antigens, as shown in Supplementary Table 1. For example, residue number 57 in Pertuzumab is Serine, but Alanine, Serine, Threonine, and Valine were found in this position in binders of all three antigens – canonical HER2, S310Y, and S310F. Subsequently, sequences composed of the residues detailed in Supplementary Table 1 in the fourteen tested positions were taken for further analysis as monoclones. This resulted in two tri-specific candidate sequences, shown in Figure 6B. These sequences were synthesized and inserted into yeast to assess binding.

### FACS Results of Chosen Colonies Showed Multi-specific Binding

FACS analysis for yeast clones expressing each of the five chosen variants from Figure 6A and 3B showed that all of them are multi-specific and detectably bind all three variants of HER2 (Supplementary Figure 3). Clone 32, however, showed weak binding to the S310F mutant; thus, we eliminated it from further analysis and took the other four clones for evaluation as IgGs.

### ELISA of Five IgGs Shows Multi-specificity for all Antigens

IgGs of the four best clones were produced and tested for binding of all three antigens, as shown in Figure 6C-E. Kd values were calculated and are shown in Figure 6F. All antibodies bound all three variants at a single-digit nM affinity.

### Engineered IgGs Have Similar T_m_ to Pertuzumab

IgGs 85 and 41 showed better antigen binding based on Supplementary Figure 4, thus, they were chosen for further analysis. Protein stability and T_m_ were determined for Pertuzumab and both IgGs using a thermal shift assay. Supplementary Figure 4A shows that the unfolding rates based on temperature for the three IgGs are very similar. Supplementary Figure 4B shows the T_m_ values for the three IgGs. These results show that the engineered IgGs maintain the original stability of Pertuzumab.

### A Cell-Based Assay Shows High HER3-Mutant HER2 Dimerization

A cell-based assay was designed to assess the functional effect of the antibodies. The assay measures signal transduction leading to HER3 phosphorylation as a result of HER2-HER3 dimerization in the presence of HRG ligand and whether a given antibody inhibits it. An initial measurement of HER3 phosphorylation is shown in Supplementary Figure 5. HER3 phosphorylation is indicative of HER2-HER3 dimerization levels. The HER2 mutant assays showed higher phosphorylated HER3 levels, implying that HER2 mutants increase HER2-HER3 dimerization levels, confirming previous findings [41, 43].

### Engineered IgGs Lower HER3 Phosphorylation

Figure 7A shows a schematic representation of HER2-HER3 dimerization. Pertuzumab disrupts the dimerization complex and abrogates tyrosine phosphorylation. Thus, we expected to see lower HER3 phosphorylation for canonical and mutant HER2 binders. The response was measured in the cell-based assay for antibody concentrations of 1nM and 100nM. Figure 7B-D show the effect of each tested IgG on HER3 phosphorylation in the presence of each: canonical HER2, the S310Y, or S310F. Pertuzumab did not inhibit the S310Y and S310F mutants, as expected. Engineered IgG 85 showed inhibition of canonical HER2, and partial inhibition of the S310F mutant. At 100nM, engineered IgG 41 completely eliminated HER3 phosphorylation for all HER2 variants, inhibiting canonical HER2 as well as both S310 mutants.

**Figure 7.**
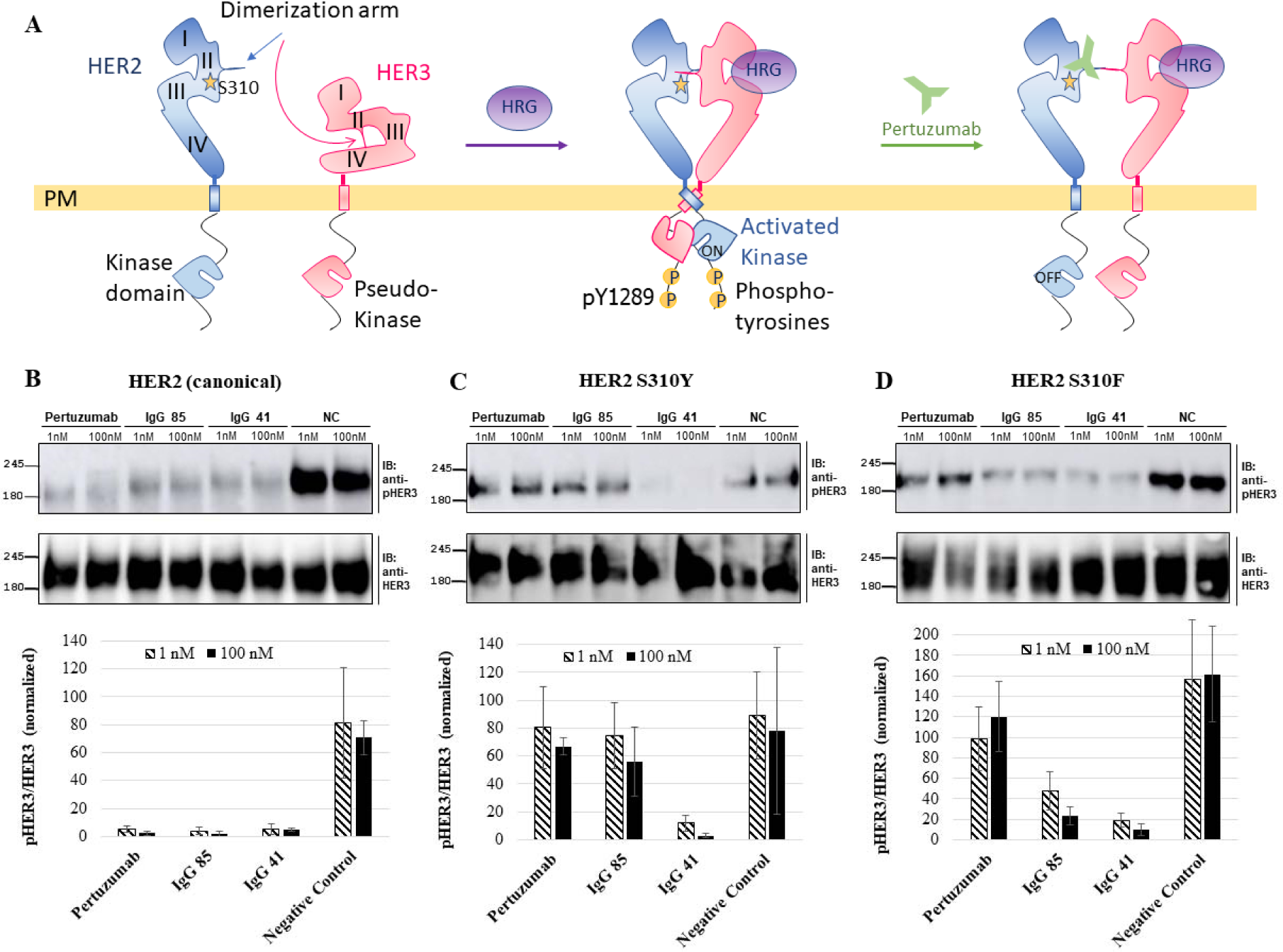
**A** Schematic representation of HER2 and HER3. HER2 S310 is represented by a yellow star in domain II. HRG ligand (purple) promotes HER2-HER3 dimerization and trans-phosphorylation, Pertuzumab (green) disrupts the dimerization complex and abrogates tyrosine phosphorylation. **B-D** HER3 phosphorylation (pHER3) due to HER2-HER3 dimerization, in antibody concentrations of 1 and 100nM for Pertuzumab and the two engineered IgG85 and IgG41. Different HER2 variants were used for dimerization: **B** canonical HER2 **C** S310Y mutant **D** S310F mutant. Quantification is based on cross-referenced western blot intensity densitometry from the mean ± SEM of three repeats. While phosphorylation was restored for the two HER2 S310 mutants in the presence of Pertuzumab, IgG 85 showed lower phosphorylation for S310F, and IgG 41 inhibited both HER2 mutants as well as canonical HER2.

## Discussion

Naturally occurring genomic variation can cause altered drug responses, ranging from improved binding to complete drug resistance [49]. Multi-specific antibodies may offer a potential solution for variations that impair drug binding, with a single therapeutic antibody that binds several versions of the target with high affinity and specificity. Here, we show a computationally-aided design framework for designing multi-specific binders based on an existing binder of a single variant.

While the antibodies that emerged from our library may require additional refinements to become clinical leads, the results highlight two important facts. First: only five mutations sufficed to add new specificities to a mature antibody. This illustrates the plasticity of mature antibodies and the fact that they can often be further engineered to acquire an additional specificity. For at least two of our variants, this new specificity comes without paying a cost in the affinity to the original target. This may suggest that there is more room for improvement either in terms of improved affinity or in terms of additional specificities that could be added with more carefully selected mutations. Second, the simple computational design that focused on rationally identifying a handful of positions to alter in the antibody to accommodate the new specificities allowed for exhaustive screening of a tiny fraction of the sequence space. This, we proposed, is crucial when attempting to introduce new specificities to mature antibodies.

HER2, like many other targets, harbors point mutations that may occur somatically in cancer but are also prevalent as germline alleles in the population. In such cases, multi-specific Abs that efficiently bind multiple variants may be beneficial both by allowing efficacy on a wider initial patient population and by increasing the therapeutic index, as it blocks at least some of the tumor’s escape routes from the drug. Thinking about these mutations early, before nominating a lead candidate for development, may save the effort, cost, and time of developing a new, mutant-specific version of the drug later. Resistance mutations prior to or as a result of tyrosine kinase inhibitors (TKIs) therapy have been targeted using second and even third-generation drugs, that are specific to different sets of mutations [50]. However, these are small molecules that do not possess the inherent plasticity that an antibody has due to its size. The results presented here suggest that such a scenario can be preempted to a certain degree by analyzing the patient population and considering variations early.

In a wider context, approaches like the one we propose here offer a path for dealing with one of the great challenges of personalized medicine, *i.e.*, designing a single molecule that is made the same for everyone but can act on multiple different stratified populations. The drug is not personalized yet its activity may be.

To identify the mutations, we used previously sequenced cancer and germline samples, enabling the use of the existing wealth of data to find mutations that can affect drug binding. Whether other mutations located in drug-target interfaces disrupt drug binding still remains to be discovered. However, the approach we described here, may allow for further engineering to cover additional escape mutations, if such mutations are identified.

## Materials and Methods

### Mutations Dataset

Somatic cancer mutations in genes were downloaded from COSMIC[3]. The COSMIC Mutation Data (Targeted Screens and Genome Screens) dataset was chosen, Release v85, 8^th^ May 2018. Records that lacked information *e.g.*, unknown mutation description or amino-acid data were removed.

### Antibodies Dataset

Therapeutic antibodies and their solved structures were acquired from Thera-SabDab[51]. Structures that do not contain the antigen (unbound) were removed. This list was crossed with the FDA-approved antibodies list. The relevant structures were downloaded from RCSB PDB[52] (http://www.rcsb.org/).

### Identifying Genes’ Locations of the Antigens

Antigens sequences were taken from the PDB files and were aligned against the hg38 genome using BLAT[53]. Antigens that were found to be of non-human origin were removed (along with their Ab).

### Obtaining Transcripts Sequences for Mutation Data

Since the residue numbering in the PDB and the mutation data are not the same, all mRNA sequences for all antigens were downloaded from GenBank[54]. From these sequences, only transcripts that contained the same numbering as the mutated data were taken (*e.g.*, if the mutation data indicated reference Leucine in position X for a specific protein, only transcripts with Leucine in that position were taken).

### Aligning the Transcripts Against PDB Sequences

To find the mutated residues in the PDB file, the mRNA sequence was first aligned against the PDB SEQRES sequence using Needle[55] global alignment tool. Since the PDB ATOM data may lack some residues that were not crystallized, the PDB ATOM residues were converted into sequences and aligned against the PDB SEQRES using Needle. Using these alignments, a script was written to find the alignment between the mRNA sequences to the PDB ATOM data.

### Identifying Contact Residues

All distances between the Ab and antigen atoms were calculated. An antigen atom was considered to be in contact if at least one of the Ab atoms was in a distance of less than 5Å.

### Mutations Filtering

Gene names obtained from the BLAT search (with all possible name variations) were used to filter COSMIC mutation data. Then, using the alignment between the mRNA sequences and the PDB ATOM data, only mutations on the binding site were taken.

### Mutated Targets Stability Prediction

The mutated target 1S78 (HER2) stability was examined using Maestro[46]. The protein structure was preprocessed (at 6.5≤pH≤7.5) and optimized to correct possible errors. The stability and affinity for each mutation (S310F, S310Y) were calculated, examining chain B, which contains the mutation, against all the other chains. S310 is numbered S288 on the PDB structure.

### Library Design

A library of sequences was designed to find an antibody that will bind both the canonical SER310 HER2 and the mutations S310Y and S310F. As only Pertuzumab heavy chain was found to be in contact with S310, its residues that are either in direct contact with SER310, adjacent to a residue that is in direct contact with S310, or in contact with HER2 and can potentially strengthen Ab-Ag binding were collected. All the bonds that these residues were involved in were calculated, as well as their evolutionary profile. Out of all collected residues, fourteen were shown to be evolutionary diverse between a hundred homologous sequences and were not involved in bonds that are likely to be essential for protein folding. In Figure 6A, these fourteen residues are listed with their classifications. Figure 2 shows these residues on the 1S78 structure. Image was generated using PyMOL[56]. Creating a library that will contain all possible residues combinations with general codons in these fourteen positions will result in a complexity of 6.23×10^18^, which is too high for an efficient Yeast-Display experiment. To reduce complexity, the residues were divided into four groups based on their sequence position. For each library sequence, a predetermined number of mutations from each group was selected, as described in Figure 6C, resulting in a complexity of 3.86×10^11^ sequences. Groups 1 and 2 which contain four and six residues, respectively, had two mutations each in every library sequence. Groups 3 and 4, having two residues each, had one mutation in every library sequence.

### Protein Expression and Purification of Canonical HER2 and Mutants

Constructs of human canonical HER2 and mutants were prepared by PCR amplification from the complete cDNA clone of human HER2 (Acc. No. BC156755 / P04626, Harvard deposit plasmid ID HsCD00348391). The extracellular region of HER2 (spanning residues 22-587) was amplified and ligated into a modified p3XFLAG-CMV™-25 Expression Vector (Sigma-Aldrich, cat: E9408) containing an N-terminal FLAG tag and a C-terminal hexahistidine tag, followed by a stop codon. The HER2 mutants S310Y and S310F were generated by assembly PCR and cloned as previously described. The proteins were expressed in HEK293F expression system (Invitrogen, cat: R790-07). HEK293F cells were maintained in FreeStyleTM 293 Expression Medium (Invitrogen, cat: 12338018) in an orbital shaker incubator at 37°C, 120 rpm, 8% CO_2_. For transfection, cells were seeded at 1×10^6^ cells/ml in a final volume of 250ml and transfected using linear polyethyleneimine (PEI)(Polysciences, Inc, Polyethylenimine HCL MAX, MW 40000, cat: 24765-1), 3µg PEI for 1µg plasmid DNA for 1×10^6^ cells. Individual cultures were grown for 7 days post-transfection and then harvested at 1500xg for 20 min at 4LJC. The supernatant was filtered and loaded onto a metal-chelate column (HisTrap, GE Healthcare) pre-equilibrated with buffer A (50mM phosphate buffer, pH 8, 0.4 M NaCl, 5% glycerol) at a flow rate of 2 ml/min. The column was washed with buffer A until a stable baseline was achieved. The proteins were eluted with 150mM imidazole, and the protein-containing fractions were selected for further purification by size exclusion chromatography. The proteins were loaded on Superdex200 10/300 (GE Healthcare, cat: 17-5175-01) that was previously equilibrated with PBS. The purified proteins were split into aliquots and flash-frozen in liquid N_2_ for further binding assays.

### Library Generation

The Pertuzumab sequence was collected from PDB 1S78 [57]. The predicted scFv sequence was obtained from IGMT/DomainGapAlign [58, 59] and synthesized by Bio-Basic Custom Gene Synthesis. The 1S78 scFv library was constructed based on the AAL160 template antibody by overlapping extension PCR with degenerate oligonucleotides. PCR used to introduce diversity with NNS codons was done using Phusion high fidelity DNA polymerase (New England Biolabs USA, cat: M0530) according to manufacturer instructions in a 3-step reaction (98°C for 30LJsec, 65°C for 20LJsec, 72°C for 30LJsec, 30 cycles). Subsequently, the DNA fragments were gel-purified and assembled in equimolar ratios in a 3-step PCR reaction, as detailed above, but in the absence of primers. The assembled scFv library was amplified using forward and reverse primers adding the yeast surface display (YSD) expression vector homology recombination sequences at the 5’ and 3’ to the scFv library allowing efficient homology recombination into EBY100 yeast strain. scFv libraries were constructed with three repeats of flexible linkers of G4S between the VH and VL. The scFv library and linearized pCTCON2 plasmid [60] were co-transformed to S. cerevisiae strain EBY100 using lithium acetate [61]. Transformed cells were selected on SDCAA plates.

### Library Selection Against Canonical HER2 and the Two Mutants

#### Magnetic Activated Cell Sorting (MACS)

Selection for binders was done essentially as described by [62]. Briefly, a culture of EBY100 cells transformed with scFv library was induced to expression by transferring to galactose-containing media (SDGAA) and incubated for 48 hours at 20°C with shaking of 200 rpm. As a first strategy for enriching the suspensions with HER2 binders, MACS was performed. In the first round, 5×10^10^ yeast cells displaying scFv were collected, washed, and incubated with 100nM canonical HER2, purified as described above. After three MACS buffer washes, the antigen-binding cells were labeled with anti-Human Flag-biotin (Miltenyi Biotec cat 130-101-569) and streptavidin microbeads (Miltenyi Biotec, cat: 130-048-101) consequently. The magnetic separation of the cells was performed using an LS-column (Miltenyi Biotec, cat: 130-042-401) that was placed in a VarioMACS separator (Miltenyi Biotec Inc., Auburn, CA, USA), as described by the manufacturer’s protocol. The positively selected cells were collected and grown in SDCAA medium at 30°C for an additional MACS round, with 200nM HER2 S310Y. The MACS procedures resulted in a library variation of 1×10^7^ cells that were then subjected to FACS.

#### Fluorescence Activated Cell Sorting (FACS)

FACS was performed with FACSAria III sorter (BD Biosciences, San Jose, CA, USA) as follows: 8×10^7^ yeast cells (in the first round, 2×10^6^ in further rounds) displaying scFv were labeled with mouse monoclonal Anti-c-myc-FITC IgG (Miltenyi Biotec cat 130-116-485) and incubated with Flag-tagged canonical HER2 and the previously purified mutant proteins at room temperature for 2 hours, followed by incubation with a mouse anti-Flag IgG [5A8E5], conjugated with iFluor 647 (cat: A01811-100, Genscript). Yeast clones exhibiting higher iFluor 647 fluorescence signal relative to FITC fluorescence were screened using a FACSAria III sorter (BD Biosciences, San Jose, CA, USA), and the collected cells were pooled down to SDCAA medium for the next round of FACS screening. The library resulted from the last round was plated onto SDCAA plates and individual colonies were picked for DNA sequencing. The complete selection scheme can be seen in Figure 4.

### Protein Expression and Purification of IgGs

The 1S78 and mutant DNA templates were collected from the selected colonies of the scFv library. The light chain (LC) and the heavy chain (HC) were amplified and cloned into pSF-kappa light and pSF-heavy plasmids, respectively, by using Golden-gate protocol. The 1S78 IgGs were expressed in HEK293F expression system (Invitrogen, cat: R790-07), as described previously. For proper IgG folding and assembly, the co-transfection ratio was 1:1 LC/HC. The culture was grown for 6 days post-transfection and then harvested at 1500xg for 20 min at 4LJC. The clear supernatant was dialyzed against 2L of phosphate-buffered saline (PBS), pH 7.4, and then loaded on a protein A column (Monofinity A resin, GenScript, cat: L00433). The elution was performed with 0.1M glycine-HCl pH 3 and directly neutralized with a 10% volume of 1M Tris–HCl, pH 8.5. One of the eluted protein fractions was loaded on Superdex200 10/300 (GE Healthcare, cat: 17-5175-01) that was previously equilibrated with PBS. The purified protein was flash-frozen in liquid N_2_ for further assays.

### ELISA

For ELISA, Microlon, High binding flat-bottomed 96-well plates were coated with 500 ng of either 1S78 IgG WT or mutants in 50mM bicarbonate buffer pH=9.6 (50 µL per well) and incubated overnight at 4°C. After three washes with PBS, plates were blocked (PBS + 3% milk) for 2 hours at RT with gentle shaking. The antigens (canonical HER2, S310Y, and S310F) were prepared at 1µM in a blocking buffer and transferred to the corresponding rows in duplicate. Plates were incubated for 60 min at RT with gentle shaking and washed subsequently. Plates were incubated with 50 µL/well of anti-Flag HRP (1:4000 diluted in blocking buffer) for 60 min at RT. A final wash step was performed and plates were developed using TMB reagent (SouthernBiotechTM) and 0.1M HCl stop solution. The optical density (OD) at 450nm was read on the Infinite 200 Pro multimode plate reader (Tecan Group Ltd., Switzerland).

### Thermal Shift Assay for Protein Stability Measurement

All measurements were performed on a CFX96 Real-Time System (C1000 Touch Thermal Cycler, Bio-Rad, Hercules, CA, USA). Each reaction was at a final volume of 25 µL, containing 5 µL of SYPRO Orange 100x (Sigma-Aldrich, cat: S5692) and protein at a final concentration of 1µM in PBS. All compounds were mixed on ice in an 8-well PCR tube. Fluorescence was measured from 4°C to 80°C with 0.5°C/10 sec steps in duplicates for each protein. The fluorescent signals that were obtained from the Texas Red channel were analyzed as follows: The replicates were normalized to a two-state model by a transition to percent values and then fitted for T_m_ calculation.

### Cell Lines

HEK293F cells (Thermo Fisher Scientific) are maintained in a FreeStyleTM 293 expression medium (Invitrogen, cat: 12338018) in suspension or adherent culture at 37°C, 8% CO_2_. COS-7 (ATCC) adherent cells are maintained in Dulbecco’s Modified Eagle’s Medium (DMEM) supplemented with 10% fetal bovine serum (FBS), 2 mM l-glutamine, 100 units/mL penicillin, and 100 mg/ml streptomycin, at 37°C, under 5% CO_2_.

### Cell-based Assay

Full-length HER2 (canonical, S310Y, and S310F) and full-length HER3 constructs were prepared by PCR amplification from the complete human cDNA clones (Acc. No. BC156755 and Acc. No. NG_011529, respectively). The genes were amplified and ligated into a modified p3XFLAG-CMV™-25 Expression Vector (Sigma-Aldrich, cat: E9408) containing a C-terminal hexahistidine tag for all, and an N-terminal FLAG tag for HER2 clones only. The mutagenesis was performed as previously described.

For transfection, COS7 cells were seeded at 1.5×10^5^ cells per well in a 6-well plate, cultured for 24h, and transiently transfected with 5µg DNA using linear polyethyleneimine (PEI) (Polysciences, Inc, Polyethylenimine HCL MAX, MW 40000, cat: 24765-1), 15 µl PEI for 5 µg plasmid DNA per well. Transfected cells were incubated at 37°C, 5% CO_2_ for 8h. The medium was then changed to a serum-free medium for starvation. Pertuzumab IgG, Negative control IgG, and our engineered 41 and 85 IgGs were added at several concentrations for 16h. The cells were then stimulated with 10nM NRG1β (PeproTech, cat: 100-03) for 10 min at 37 °C, cooled on ice for 5 min, washed two times in ice-cold PBS, and lysed in 400 µl SIMA buffer (120mM NaCl, 25mM HEPES, 1% Triton, 10% glycerol, 1 mM EGTA, 0.75mM MgCl2, Roche Complete protease inhibitors, 1mM sodium orthovanadate,2 mM sodium fluoride, pH 7.4) per 6-well on ice for 30 min. Lysates were transferred into 1.5ml microcentrifuge tubes, spun at 15,000g for 15 min, and supernatants were transferred into fresh tubes with SDS loading dye. Phospho-HER3 (pY1289) and HER3 levels in lysates were determined by western blot. Antibodies used for detection were rabbit anti-HER3 (Cell Signaling, D22C5, 1:1,000), rabbit anti-phospho-HER3 recognizing phosphorylated tyrosine position Y1289 (Cell Signaling, 21D3, 1:1,000) and anti-rabbit IgG HRP-linked antibody (Cell Signaling, cat: 7074, 1:10,000). Protein visualization by chemiluminescence and quantification was performed using AmershamTM Imager 600 (GE Healthcare).

## Acknowledgments

We thank Michael Zhenin (Biolojic Design Ltd.) for mutations assessment and library design assistance, Shmuel Bernstein (Biolojic Design Ltd.), and Tal Vana (Biolojic Design Ltd.) for assistance with the experiments.

## Author contributions

Y.O. conceived the study. Y.O., S.P., J.G.H., G.N., S.F., and Y.F. designed the experiments. S.P., J.G.H., and N.Z.B. performed the experiments. Y.O., S.P., and J.G.H. analyzed the results and wrote the paper.

