## Supplementary Figures for "Computational Engineering of a Therapeutic Antibody to Inhibit Multiple Mutants of HER2 Without Compromising Inhibition of the Canonical HER2"

```
1S78_B             1 ----------------------TQVCTGTDMKLRLPASPETHLDMLRHLY     28
                                           ||||||||||||||||||||||||||||
P04626             1 MELAALCRWGLLLALLPPGAASTQVCTGTDMKLRLPASPETHLDMLRHLY     50

1S78_B            29 QGCQVVQGNLELTYLPTNASLSFLQDIQEVQGYVLIAHNQVRQVPLQRLR     78
                     ||||||||||||||||||||||||||||||||||||||||||||||||||
P04626            51 QGCQVVQGNLELTYLPTNASLSFLQDIQEVQGYVLIAHNQVRQVPLQRLR    100

1S78_B            79 IVRGTQLFEDNYALAVLDNGDPL---------SPGGLRELQLRSLTEILK    119
                     |||||||||||||||||||||||         ||||||||||||||||||
P04626           101 IVRGTQLFEDNYALAVLDNGDPLNNTTPVTGASPGGLRELQLRSLTEILK    150

1S78_B           120 GGVLIQRNPQLCYQDTILWKDIFHKNNQLALTLIDTNRSRACHPCSPMCK    169
                     ||||||||||||||||||||||||||||||||||||||||||||||||||
P04626           151 GGVLIQRNPQLCYQDTILWKDIFHKNNQLALTLIDTNRSRACHPCSPMCK    200

1S78_B           170 GSRCWGESSEDCQSLTRTVCAGGCARCKGPLPTDCCHEQCAAGCTGPKHS    219
                     ||||||||||||||||||||||||||||||||||||||||||||||||||
P04626           201 GSRCWGESSEDCQSLTRTVCAGGCARCKGPLPTDCCHEQCAAGCTGPKHS    250

1S78_B           220 DCLACLHFNHSGICELHCPALVTYNTDTFESMPNPEGRYTFGASCVTACP    269
                     ||||||||||||||||||||||||||||||||||||||||||||||||||
P04626           251 DCLACLHFNHSGICELHCPALVTYNTDTFESMPNPEGRYTFGASCVTACP    300

1S78_B           270 YNYLSTDVGSCTLVCPLHNQEVTAEDGTQRCEKCSKPCARVCYGLGMEHL    319
                     ||||||||||||||||||||||||||||||||||||||||||||||||||
P04626           301 YNYLSTDVGSCTLVCPLHNQEVTAEDGTQRCEKCSKPCARVCYGLGMEHL    350

1S78_B           320 REVRAVTSANIQEFAGCKKIFGSLAFLPESFDGDPASNTAPLQPEQLQVF    369
                     ||||||||||||||||||||||||||||||||||||||||||||||||||
P04626           351 REVRAVTSANIQEFAGCKKIFGSLAFLPESFDGDPASNTAPLQPEQLQVF    400

1S78_B           370 ETLEEITGYLYISAWPDSLPDLSVFQNLQVIRGRILHNGAYSLTLQGLGI    419
                     ||||||||||||||||||||||||||||||||||||||||||||||||||
P04626           401 ETLEEITGYLYISAWPDSLPDLSVFQNLQVIRGRILHNGAYSLTLQGLGI    450

1S78_B           420 SWLGLRSLRELGSGLALIHHNTHLCFVHTVPWDQLFRNPHQALLHTANRP    469
                     ||||||||||||||||||||||||||||||||||||||||||||||||||
P04626           451 SWLGLRSLRELGSGLALIHHNTHLCFVHTVPWDQLFRNPHQALLHTANRP    500

1S78_B           470 EDECVGEGLACHQLCARGHCWGPGPTQCVNCSQFLRGQECVEECRVLQGL    519
                     ||||||||||||||||||||||||||||||||||||||||||||||||||
P04626           501 EDECVGEGLACHQLCARGHCWGPGPTQCVNCSQFLRGQECVEECRVLQGL    550

1S78_B           520 PREYVNARHCLPCHPECQPQNGSVTCFGPEADQCVACAHYKDPPFCVAR-    568
                     |||||||||||||||||||||||||||||||||||||||||||||||||
P04626           551 PREYVNARHCLPCHPECQPQNGSVTCFGPEADQCVACAHYKDPPFCVARC    600

1S78_B           569 --------------------------------------------------    568

P04626           601 PSGVKPDLSYMPIWKFPDEEGACQPCPINCTHSCVDLDDKGCPAEQRASP    650

1S78_B           569 --------------------------------------------------    568

P04626           651 LTSIISAVVGILLVVVLGVVFGILIKRRQQKIRKYTMRRLLQETELVEPL    700

1S78_B           569 --------------------------------------------------    568

P04626           701 TPSGAMPNQAQMRILKETELRKVKVLGSGAFGTVYKGIWIPDGENVKIPV    750

1S78_B           569 --------------------------------------------------    568

P04626           751 AIKVLRENTSPKANKEILDEAYVMAGVGSPYVSRLLGICLTSTVQLVTQL    800

1S78_B           569 --------------------------------------------------    568

P04626           801 MPYGCLLDHVRENRGRLGSQDLLNWCMQIAKGMSYLEDVRLVHRDLAARN    850

1S78_B           569 --------------------------------------------------    568

P04626           851 VLVKSPNHVKITDFGLARLLDIDETEYHADGGKVPIKWMALESILRRRFT    900

1S78_B           569 --------------------------------------------------    568

P04626           901 HQSDVWSYGVTVWELMTFGAKPYDGIPAREIPDLLEKGERLPQPPICTID    950

1S78_B           569 --------------------------------------------------    568

P04626           951 VYMIMVKCWMIDSECRPRFRELVSEFSRMARDPQRFVVIQNEDLGPASPL   1000

1S78_B           569 --------------------------------------------------    568

P04626          1001 DSTFYRSLLEDDDMGDLVDAEEYLVPQQGFFCPDPAPGAGGMVHHRHRSS   1050

1S78_B           569 --------------------------------------------------    568

P04626          1051 STRSGGGDLTLGLEPSEEEAPRSPLAPSEGAGSDVFDGDLGMGAAKGLQS   1100

1S78_B           569 --------------------------------------------------    568

P04626          1101 LPTHDPSPLQRYSEDPTVPLPSETDGYVAPLTCSPQPEYVNQPDVRPQPP   1150

1S78_B           569 --------------------------------------------------    568

P04626          1151 SPREGPLPAARPAGATLERPKTLSPGKNGVVKDVFAFGGAVENPEYLTPQ   1200

1S78_B           569 --------------------------------------------------    568

P04626          1201 GGAAPQPHPPPAFSPAFDNLYYWDQDPPERGAPPSTFKGTPTAENPEYLG   1250

1S78_B           569 -----    568

P04626          1251 LDVPV   1255
```

**Supplementary Figure 1.** Alignment of canonical human HER2 sequence from UniProt (P04626) and the crystalized residues of HER2 from PDB 1S78, chain B. Alignment was done using EMBOSS Needle. S310 from the original sequence / S288 from the 1S78 can be seen on the seventh block of the alignment and are highlighted in yellow.

>1

TQDQNSGSIQVAGS

>2

TQDQNSGSQQVEGW

>3

TEDTNSGSIQVAGW

>4

SNDTNSGAIQVAGW

>5

SQDTNSGVIQVVGL

>6

SPDTNSGTIQVSGS

>7

SQDTNSGSIHVAGL

>8

SQDTNSGMIQVEGS

>9

NWDTNSGIIQVGGS

>10

QRDTNSGSIRVEGS

>11

SMDTNSGSLAVEGL

>12

TQDTNSGSVQVEGW

>13

SPDTNSGVIRVRGY

>14

SQDTNSGGIRVSGS

>15

TQDTNSGLIRVLGS

>16

GQDTNSGSIQVAGS

>17

ETSTNSGVIQVPGS

>18

TQDTNSGSIRVRGW

>19

SHDTNSGSIQVPGS

>20

SPDTNSGSIQVRGS

>21

SQDTNSGSIPVEGM

>22

GQDTNSGSIQVEGS

>23

TLDTNSGVIRVEGS

>24

GQDTNSGSIQVEGS

>25

SFDTNSGVIQVPGN

>26

GQDTNSGVIQVRGW

>27

EQDTNSGVIQVGGW

>28

TRDTNSGAIQVEGW

>29

GYDTNSGVIQVPGS

>30

TGETNSGSIKVRGW

>31

TGLTNSGTIQVPGL

>32

TGETNSGSIKVRGW

>33

TGLTNSGVIQVSGN

>34

TLSTNSGVIQVVGS

>35

TGLTNSGCIKVAGS

>36

TQDTNSGVIQVPGW

>37

TLSTNSGVVQVEGL

>38

TMATNSGAIQVEGL

>39

TGLTNSGAIQVTGW

>40

TGLTNSGAIPVWGL

>41

TWSTNSGVIKVGGS

>42

TPLTNSGVIQVAGS

>43

TGLTNSGTIQVEGW

>44

TSSTNSGVIQVRGW

>45

TLATNSGVIQVEGH

>46

TGLTNSGTIRSRGM

>47

TRATNSGTIEVEGS

>48

TGLTNSGAIQVNGS

>49

TRSTNSGSISVEGL

>50

TPLTNSGAIQVAGS

>51

TWDTNSGVIQVGGS

>52

TQSTNSGAINVEGF

>53

TGLTNSGVIQVAGS

>54

TPLTNSGSITVAGN

>56

TNSTNSGVIQVAGS

>57

TRATNSGALQVEGW

>58

TRATNSGVISVEGS

>59

TDLTNSGVIQVRGS

>60

TRSTNSGTIQVNGS

>61

TQATNSGAIQVEGS

>62

TRSTNSGAIPVNGF

>63

TRLTNSGAIQVEGN

>64

TLSTNSGVISVNGS

>65

TGLTNSGVIKVWGK

>66

TGLTNSGAIQARGS

>67

TGLTNSGGIRVRGS

>68

TGLTNSGSISVRGW

>69

TRATNSGAIKVEGN

>70

TQMTNSGVIQVPGS

>71

TGLTNSGAIRVWGN

>72

TGLTNSGVIQVRGS

>73

TGLTNSGSIPSRGW

>74

TGLTNSGAINVSGS

>75

TGLTNSGTLQVAGW

>76

TPLTNSGVIQVGGS

>77

TGLTNSGAIQSRGS

>78

TWATNSGVITVGGS

>79

TRSTNSGVIQVEGF

>80

TGLTNSGAIQVGGS

>81

TGLTNSGVIRSRGS

>82

TMATNSGAIQVEGL

>83

TLATNSGVIQVPGM

>84

TQLTNSGTIRVEGS

>85

TRSTNSGVIQVEGS

>86

TRATNSGSIPVEGL

>87

TGLTNSGVIQVRGS

>88

TPLTNSGVIQVRGS

>89

TGLTNSGVIQVTGM

>90

TPLTNSGAIQVAGS

**Supplementary Figure 2.** The 14 positions that were mutated (see Figure 3A), from the sequences of the 90 colonies from the fourth round of selection (R4). Colonies 1-30 were taken from the canonical HER2 binding population (Figure 5B-II,VI), colonies 31-60 were taken from S310Y HER2 binding population (Figure 5B-III,VII) and colonies 61-90 were taken from S310F HER2 binding population (Figure 5B-IV,VIII).

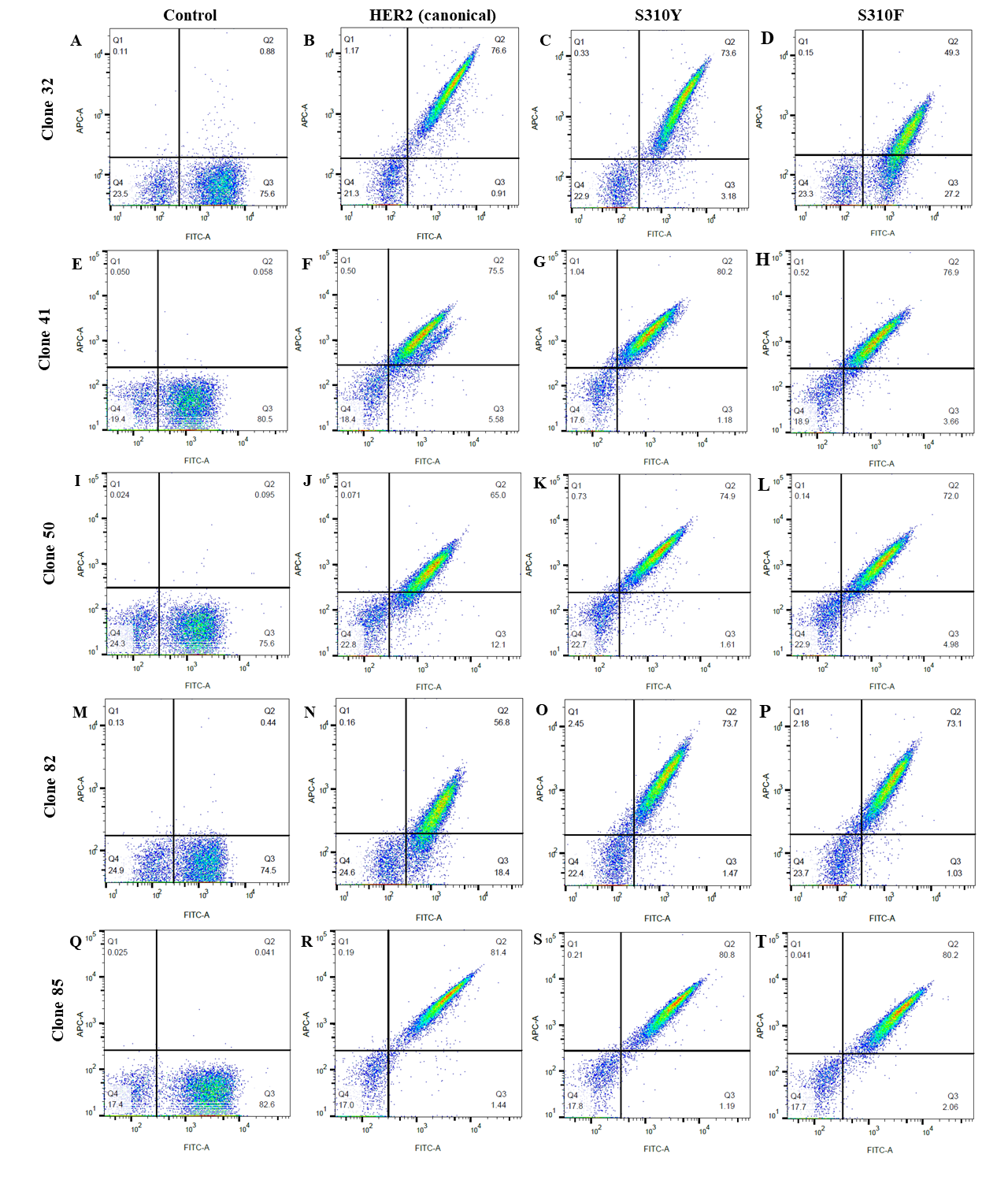

**Supplementary Figure 3.** FACS results for all five clones tested for multi-specificity. Each row shows the results of each clone: **A-D** clone 32 **E-H** clone 41 **I-L** clone 50 **M-P** clone 82 **Q-T** clone 85. First column shows control - with no antigen, second to fourth columns are FACS results against canonical HER2, S310Y and S310F mutants, respectively.

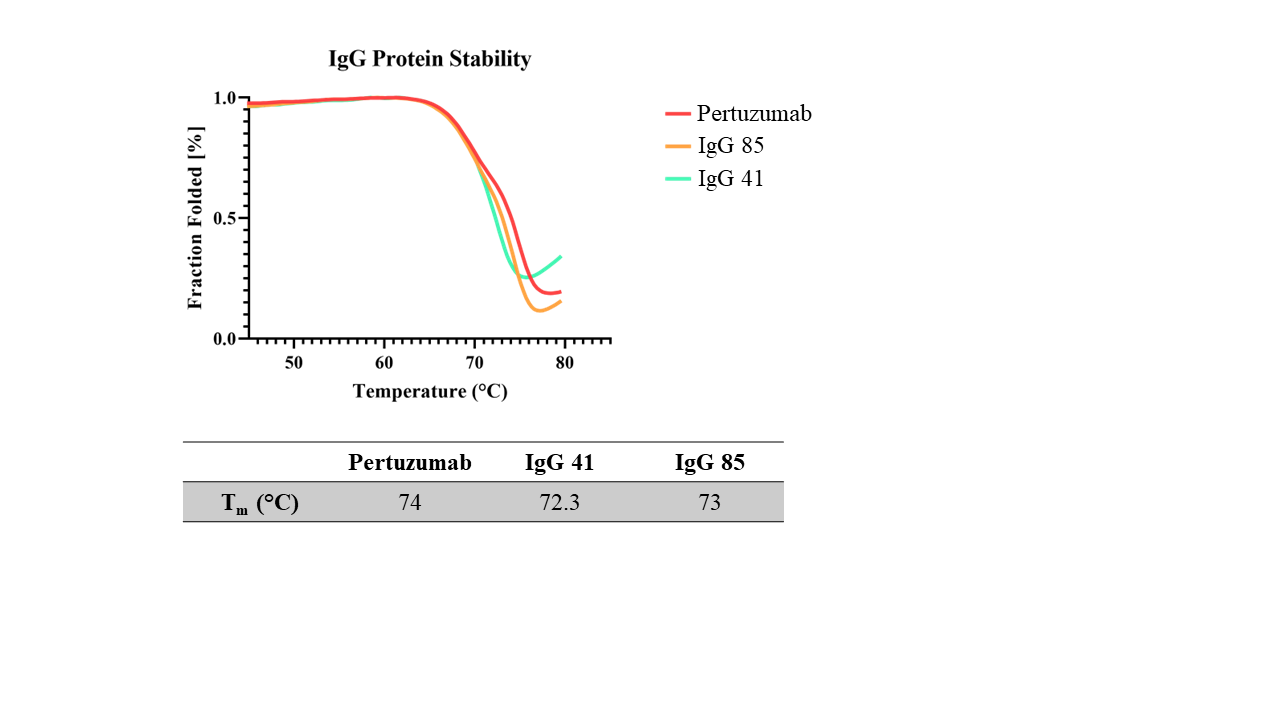

**A**

**B**

**Supplementary Figure 4.** Thermal shift assay results. **A** Unfolding rates based on temperature. **B** T_m_ values for each IgG. The two engineered IgGs are highly stable and show similar thermal-induced unfolding patterns and T_m_s to canonical Pertuzumab.

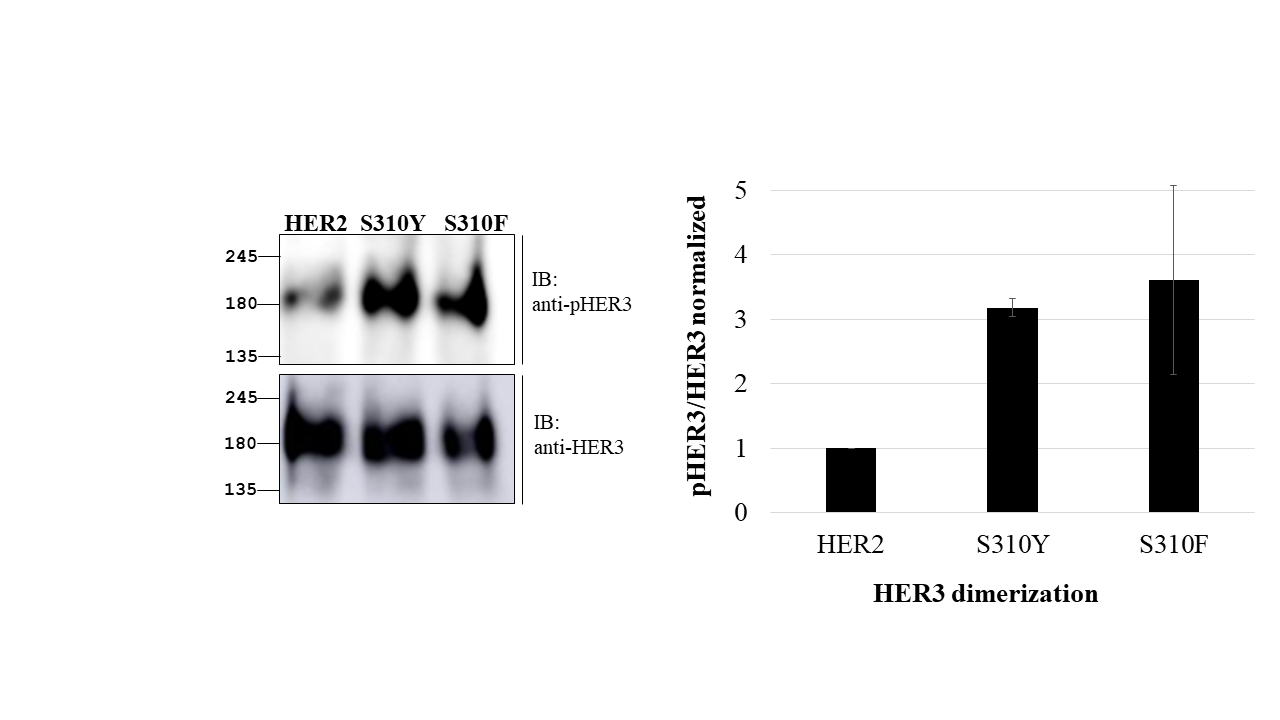

**A**

**B**

**Supplementary Figure 5.** HER3 phosphorylation (pHER3) caused by HER2-HER3 dimerization, with each of the HER2 variants: canonical HER2, S310Y mutant, and S310F mutant. **A** Western blots. Top photo: phosphorylated HER3. Bottom photo: HER3. **B** Quantification, based on cross-referenced western blot intensity densitometry from the mean ± SEM of three repeats. HER2 mutants show higher HER3 phosphorylation, indicating an increase in HER2-HER3 dimerization.

**Supplementary Tables**

| Position | Potential Residues | | | | | | |
| --- | --- | --- | --- | --- | --- | --- | --- |
| THR28 | T |  |  |  |  |  |  |
| THR30 | G | M | L | Q | P | R | W |
| ASP31 | S |  |  |  |  |  |  |
| THR33 | T |  |  |  |  |  |  |
| ASN53 | N |  |  |  |  |  |  |
| SER54 | S |  |  |  |  |  |  |
| GLY55 | G |  |  |  |  |  |  |
| SER57 | A | S | T | V |  |  |  |
| ILE58 | I | L |  |  |  |  |  |
| GLN61 | K | Q | P | R |  |  |  |
| VAL71 | V |  |  |  |  |  |  |
| ARG73 | A | E | G | P | S | R |  |
| GLY97 | G |  |  |  |  |  |  |
| SER99B | M | L | N | S | W |  |  |

**Supplementary Table 1.** The fourteen chosen residues from Figure 1A. For each position, residues that are present in binders of all three antigens (canonical HER2, S310Y, and S310F), are listed.
